# Human induced pluripotent stem cell-derived ovarian support cell co-culture improves oocyte maturation *in vitro* after abbreviated gonadotropin stimulation

**DOI:** 10.1101/2023.03.27.534479

**Authors:** Sabrina Piechota, Maria Marchante, Alexa Giovannini, Bruna Paulsen, Kathryn S Potts, Graham Rockwell, Caroline Aschenberger, Alexander D Noblett, Alexandra B Figueroa, Marta Sanchez, Ferran Barrachina, Klaus Wiemer, Luis Guzman, Pedro Belchin, Merrick Pierson Smela, Patrick R.J. Fortuna, Pranam Chatterjee, Nam D Tran, Dawn A Kelk, Marcy Forti, Shelby Marcinyshyn, Trozalla Smith, David H McCulloh, Marta-Julia Fernandez-Gonzalez, Silvia Ortiz, Joshua U Klein, Peter Klatsky, Daniel Ordonez-Perez, Christian C Kramme

## Abstract

Assisted reproductive technologies (ART) have significantly impacted fertility treatment worldwide through innovations such as *in vitro* fertilization (IVF) and *in vitro* maturation (IVM). IVM holds promise as a technology for fertility treatment in women who cannot or do not wish to undergo conventional controlled ovarian stimulation (COS). However, IVM has historically shown highly variable performance in maturing oocytes and generating oocytes with strong developmental capacity. Furthermore, recently reported novel IVM approaches are limited to use in cycles lacking human chorionic gonadotropin (hCG) triggers, which is not standard practice in fertility treatment. We recently reported the development of ovarian support cells (OSCs) generated from human induced pluripotent stem cells (hiPSCs) that recapitulate dynamic ovarian function *in vitro*. Here we investigate the potential of the
se OSCs in an IVM co-culture system to improve the maturation of human cumulus-enclosed immature oocytes retrieved from abbreviated gonadotropin stimulated cycles. We reveal that OSC-IVM significantly improves maturation rates compared to existing IVM systems. Most importantly, we demonstrate that OSC-assisted IVM oocytes are capable of significantly improving euploid blastocyst formation and yielding blastocysts with normal global and germline differential methylation region methylation profiles, a key marker of their clinical utility. Together, these findings demonstrate a novel approach to IVM with broad applicability to modern ART practice.

**Structured Abstract:** *Objective:* To determine if *in vitro* maturation (IVM) of human oocytes can be improved by co-culture with ovarian support cells (OSCs) derived from human induced pluripotent stem cells (hiPSCs).

*Design:* Three independent experiments were performed in which oocyte donors were recruited to undergo abbreviated gonadotropin stimulation and retrieved cumulus oocyte complexes (COCs) were randomly allocated between the OSC-IVM and control IVM conditions.

*Subjects:* Across the three experiments, a total of 67 oocyte donors aged 19 to 37 years were recruited for retrieval using informed consent. Anti-mullerian hormone (AMH) value, antral follicle count (AFC), age, BMI, and ovarian pathology were used for inclusion and exclusion criteria.

*Intervention and Control:* The OSC-IVM culture condition was composed of 100,000 OSCs in suspension culture supplemented with human chorionic gonadotropin (hCG), recombinant follicle stimulating hormone (rFSH), androstenedione and doxycycline. IVM controls comprised commercially-available IVM media without OSCs and contained either the same supplementation as above (media-matched control), or FSH and hCG only (IVM media control). In one experiment, an additional control using fetal ovarian somatic cells (FOSCs) was used with the same cell number and media conditions as in the OSC-IVM.

*Main Outcome Measures:* Primary endpoints consisted of metaphase II (MII) formation rate and oocyte morphological quality assessment. A limited cohort of oocytes were utilized for secondary endpoints, consisting of fertilization and blastocyst formation rates with preimplantation genetic testing for aneuploidy (PGT-A) and embryo epigenetic analysis.

*Results:* OSC-IVM resulted in a statistically significant improvement in MII formation rate compared to the media-matched control, a commercially available IVM media control, and the FOSC-IVM control. Oocyte morphological quality between OSC-IVM and controls did not significantly differ. OSC-IVM displayed a trend towards improved fertilization, cleavage, and blastocyst formation. OSC-IVM showed statistically significant improvement in euploid day 5 or 6 blastocyst formation compared to the commercially available IVM media control. OSC-IVM embryos displayed similar epigenetic global and germline loci profiles compared to conventional stimulation and IVM embryos.

*Conclusion:* The novel OSC-IVM platform is an effective tool for maturation of human oocytes obtained from abbreviated gonadotropin stimulation cycles, supporting/inducing robust euploid blastocyst formation. OSC-IVM shows broad utility with different stimulation regimens, including hCG triggered and untriggered oocyte retrieval cycles, making it a highly useful tool for modern fertility treatment.

## Introduction

It is estimated that nearly 1 in 6 women in the United States and many European nations have sought fertility treatment.^1^ Assisted reproductive technologies (ART) such as *in vitro* fertilization (IVF) offer remarkable benefits for treating aspects of infertility for certain patient populations.^2,3^ However, the use of high doses of gonadotropin in stimulation regimens has resulted in uncomfortable and sometimes serious and long lasting side effects for women undergoing the process, with rising costs continuing to limit access in many nations.^4,5^ In addition, for some women for whom conventional controlled ovarian hyperstimulation is contraindicated, standard IVF practice is not feasible due to the medical risk it poses.^6–8^ Therefore, new technologies that allow for a reduction of gonadotropin stimulation in fertility treatments are important to improve patient outcomes and ART accessibility.

One option for reducing gonadotropin usage in fertility treatment is through the application of minimal or abbreviated stimulation cycles. Such cycles drastically reduce the cost and complications of IVF, while nearly completely eliminating the chance for severe complications such as ovarian hyperstimulation syndrome (OHSS).^9–11^ Depending on the protocol, abbreviated gonadotropin stimulation results in a cohort of oocytes that are mostly or all immature, including germinal vesicle containing (GV) oocytes and oocytes lacking a germinal vesicle and polar body (metaphase I, MI), while some regimens may yield a small pool of mature oocytes with a polar body (metaphase II, MII).^11–13^ In abbreviated gonadotropin cycles, *in vitro* maturation (IVM) holds promise as an ART strategy to mature oocytes outside of the body for use in fertility treatments. Numerous studies have employed IVM effectively, often through the application of reproductive cell culture media supplemented with growth factors, small molecules, and gonadotropins.^11,13–26^ However, many of these studies have shown mixed results of IVM applications in terms of oocyte maturation rate, cleaved embryo formation, and blastocyst formation with some studies showing lower pregnancy rates from IVM cycles. Therefore, improvement is needed in IVM strategies that yield robust, high-quality oocytes with strong developmental competence.

Oocyte maturation is a coordinated series of molecular processes occurring within the oocyte nucleus and cytoplasm.^27,28^ These molecular processes result in both nuclear maturation, namely extrusion of the first polar body (PB1), assembly of the second meiotic spindle, and cytoplasmic maturation, defined as distribution of organelles in the oocyte cytoplasm and deposition of proteins and transcripts needed for fertilization competence and subsequent embryogenesis.^29^ Oocyte maturation normally occurs within the ovary in response to complex, temporally-regulated, extrinsic signals provided through hormone signaling, growth factor production, and nutrient dynamics in the follicular environment.^28,30–36^ Many of these processes are regulated by somatic ovarian cells, such as granulosa, theca, and stroma cells. Limited but promising research has been performed to assess the use of these somatic cell types to support human oocyte *in vitro* maturation applications.^37–40^ However, the use of patient-derived somatic cell types besides cumulus cells is challenging due to inconsistent cell type composition retrieval, difficulty in extracting and purifying populations, and infeasibility of extracting sufficient support cells without harming patients. In addition, some patients’ infertility may arise from inappropriate signaling within and among their own follicular cells, limiting their utility for IVM co-culture. Therefore, new IVM systems that effectively and consistently deliver specific ovarian support cell types for co-culture with immature oocytes is a salient potential platform for human oocyte IVM.

Our previous work described the generation of ovarian support cells (OSCs) from human induced pluripotent stem cells (hiPSCS) in a rapid, efficient, and reproducible manner through transcription factor (TF)-directed differentiation.^41^ The OSCs primarily are composed of FOXL2+ AMHR2+ NR2F2+ granulosa-like cells. The OSCs are steroidogenic, producing aromatase in response to follicle stimulating hormone (FSH) stimulation to catalyze the estradiol synthesis pathway, and produce the necessary growth factors needed for robust paracrine interaction with oocytes and cumulus cells. In this study, we investigated the potential of hiPSC-derived OSCs to improve *in vitro* maturation of oocytes in a co-culture system with immature cumulus oocyte complexes (COCs) retrieved from subjects following abbreviated gonadotropin cycles.

## Materials and Methods

### Type of Study

The study described here is a basic science study, designed to evaluate the oocyte maturation-stimulating potential of a novel *in vitro* maturation (IVM) system and its effect on oocyte quality. No gametes or embryos utilized or created in this study were used for clinical trials or reproductive purposes such as long-term banking, transfer, implantation, or for use by gamete donors.

### Subject ages, IRB and Informed Consent

This study was performed according to the ethical principles of the Declaration of Helsinki. Oocyte donor subjects were enrolled in the study through Ruber Clinic (Madrid, Spain), Spring Fertility Clinic (New York, USA), Extend Fertility Clinic (New York, USA), and Pranor Clinic (Lima, Peru) using informed consent to donate gametes for research purposes (CNRHA 47/428973.9/22 (Spain), Western IRB # 20225832 (USA), and Protocol #GC-MSP-01 (Peru) respectively). Oocytes retrieved and donated from the Ruber and Pranor clinics were consented and utilized for research purpose for maturation analysis endpoints only, while oocytes retrieved and donated from Spring Fertility and Extend Fertility were consented and utilized for oocyte maturation and embryo formation endpoints. Sperm donor subjects were enrolled through Seattle Sperm Bank (Seattle, Washington) under informed consent through Western IRB # 20225832 (USA), for donation of gametes for research purposes involving the creation of human embryos. An additional eight euploid blastocysts were donated for research purposes for this study to serve as controlled ovarian stimulation (COS) controls for embryo epigenetic analysis under the IRB #2022-001 (ReproART, Georgia). Subject ages ranged between 19 and 37 years of age with an average of 28.

### Experimental Design

This study utilized three independent experiments, which tested hypotheses on non-overlapping patient groups at different clinics under different IRB consents and stimulation regimens. The design of these experiments is shown via a flowchart in Figure 1.

**Figure 1:**
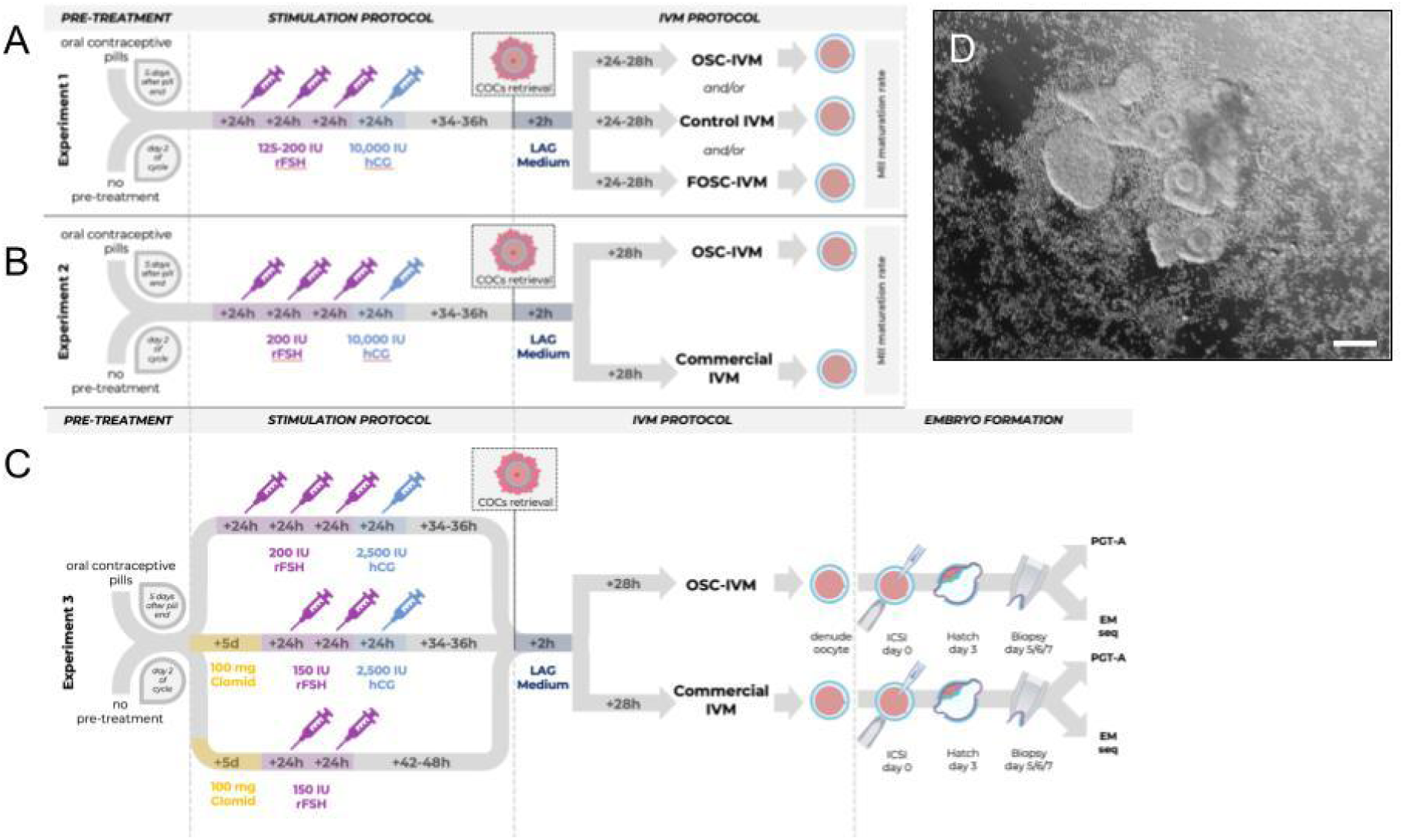
Study Design Flow Chart. Schematic representation of the three independent experiments performed in the study, depicting stimulation protocols and embryology workflows. **A** Experiment 1 refers to Figure 2. **B** Experiment 2 refers to Figure 3. **C** Experiment 3 refers to Figures 4 and 5. **D** Representative image of co-culture setup at time of plating containing human COCs (n=5) and 100,000 OSCs. Scale bar: 100μm. COCs with varying degrees of cumulus enclosure are seen with surrounding OSCs in suspension culture. COCs, cumulus-oocytes complexes; ICSI, intracytoplasmic sperm injection; IU, International Units ; rFSH, recombinant Follicle Stimulating Hormone; Clomid, clomiphene citrate; hCG, human Chorionic Gonadotropin; LAG, pre-incubation medium (MediCult); PGT-A, pre-implantation genetic testing for aneuploidy; EM-Seq; Enzymatic methylation sequencing.

#### Experiment 1 (OSC activity)

The purpose of this comparison was to determine whether the stimulated OSCs were the active ingredient of the co-culture system in a pilot study. This experiment ended with oocyte maturation analysis and involved subjects from Ruber and Pranor Clinic.

#### Experiment 2 (OSC clinical relevance)

The purpose of this experiment was to compare the efficacy of the OSC-IVM system and the commercially available *in vitro* maturation system (Medicult IVM) in maturing human oocytes. This experiment ended with oocyte maturation analysis and involved subjects from Ruber Clinic.

#### Experiment 3 (OSC clinical relevance)

The purpose of this experiment was to compare the efficacy of the OSC-IVM system and the commercially available *in vitro* maturation system (Medicult IVM) in maturing human oocytes and generating euploid blastocysts. This experiment ended with embryo formation and embryo molecular analysis and involved subjects from Spring Fertility Clinic and Extend Fertility Clinic.

### Study Duration

The period of recruitment and duration of Experiment 1 was April 1^st^, 2022 to August 1^st^, 2022. The period of recruitment and duration of Experiment 2 was October 1^st^, 2022 to December 1^st^, 2022. The period of recruitment and duration of Experiment 3 was December 1^st^, 2022 to July 1^st^, 2023.

### Subject Screening and Stimulation Characteristics

#### Experiment 1

27 subjects received three to four days of stimulation using 325-600 IU of rFSH with a 10,000 IU hCG trigger in preparation for immature oocyte aspiration. Non-contraceptive subjects began stimulation on day 2 of their menstrual cycle, while contraceptive patients began on day 5 following cessation of pills. Triggers were scheduled when 2 or more follicles reached 7-9mm in diameter, and average follicle sizes of 6-14mm were used for extraction 34 to 36 hours after trigger. Inclusion criteria included: AMH ≥ 1 ng/ml, AFC ≥ 15, ages 20-40, BMI 20-30 kg/m2, and normal ovulatory cycle. Exclusion criteria included: Diagnosis of severe endometriosis, pathology preventing access to both ovaries, medical contraindication for undergoing stimulation or anesthesia, positive test for sexually transmitted disease, clinical diagnosis of polycystic ovarian syndrome (PCOS), clinical diagnosis of hypothyroidism, and previously known defect in oocyte maturation.

#### Experiment 2

23 subjects received three consecutive days of 200 IU of rFSH with a 10,000IU hCG trigger in preparation for immature oocyte aspiration. Non-contraceptive subjects began stimulation on day 2 of their menstrual cycle, while contraceptive patients began on day 5 following cessation of pills. Triggers were scheduled when 2 or more follicles reached 7-9mm in diameter, and average follicle sizes of 6-14mm were used for extraction 34 to 36 hours after trigger. Inclusion criteria included: AMH ≥ 1.5 ng/ml, AFC ≥ 15, ages 25-35, BMI 20-30 kg/m2, and normal ovulatory cycle. Exclusion criteria included: Diagnosis of severe endometriosis, pathology preventing access to both ovaries, medical contraindication for undergoing stimulation or anesthesia, positive test for sexually transmitted disease, clinical diagnosis of PCOS, clinical diagnosis of hypothyroidism, and previously known defect in oocyte maturation.

#### Experiment 3

17 subjects received either five doses of clomiphene citrate (Clomid) (100 mg) with an additional one to two doses of 150 IU rFSH with or without a 2,500 IU hCG trigger, or three doses of 200 IU rFSH with a 2,500 IU hCG trigger. Non-contraceptive subjects began stimulation on day 2 of their menstrual cycle, while contraceptive patients began on day 5 following cessation of pills. Triggers were scheduled when 2 or more follicles reached 7-9mm in diameter, and average follicle sizes of 6-14mm were used for extraction 34 to 36 hours after trigger. In non-triggered cycles, extraction was scheduled when 2 or more follicles reached 7-9mm in diameter, and average follicle sizes of 6 to 14mm were used for extraction 42-48 hours after the last rFSH injection. Inclusion criteria included: AMH ≥ 2 ng/ml, AFC ≥ 20, ages 25-35, BMI 20-30 kg/m2, normal ovulatory cycle, TSH ≤ 3 mIU/mL. Exclusion criteria included: Diagnosis of severe endometriosis, pathology preventing access to both ovaries, medical contraindication for undergoing stimulation or anesthesia, positive test for sexually transmitted disease, clinical diagnosis of PCOS, clinical diagnosis of hypothyroidism, and previously known defect in oocyte maturation. For male gamete donors, three donors were selected ages 21-35 who were consented for research donation resulting in embryo formation, with sperm donation with the following characteristics: Normal Volume (>2.0 ml), Normal Sperm Count (concentration; >15 M/ml), Normal Motility (>40%), and Normal Morphology (>4% Strict Morphology).

A complete table of donor stimulation regimens for each donor in the study is provided in Supplementary Table 1.

### Serum hormone and follicle development monitoring during stimulation

Baseline serum evaluation of AMH levels was performed for all patients as part of inclusion criteria screening. For Experiments 1 and 2, patients were additionally monitored on the day of stimulation start, day of trigger and day of oocyte pickup (OPU). During these monitoring periods, transvaginal ultrasound was performed to assess follicle sizes and development. Additionally, serum levels of LH, E2, and P4 were measured to assess the early follicular growth phase (E2 > 100 ng/ml), to monitor for premature luteinization and for spikes in LH (LH ≥ 10 IU/ml). For Experiment 3, which was performed at a different center than Experiment 1 and 2, ultrasound monitoring and serum evaluation of E2, P4, and LH was performed at the start of stimulation and day of oocyte pick up only.

### Collection of Cumulus Oocyte Complexes (COCs)

#### Aspiration of small ovarian follicles to retrieve immature cumulus oocyte complexes

##### In Experiment 1 and 2

Aspirations were performed 34-36 hours after the trigger injection (10,000 IU hCG) using a transvaginal ultrasound with a needle guide on the probe to retrieve oocytes for co-culture experiments. Aspiration was performed using ASP medium (Vitrolife) without follicular flushing using double lumen 19-gauge needles (double lumen needles were selected due to the additional stiffness provided by the second channel inside the needle). Vacuum pump suction (100-120 mm Hg) was used to harvest follicular contents through the aspiration needle and tubing into a 15 ml round bottom polystyrene centrifuge tube.

##### In Experiment 3

For the conditions where the final outcome was embryo formation, aspirations were performed 34-36 hours after trigger injection (2,500 IU hCG) or 42-48 hours after the last rFSH injection for untriggered cycles. Aspiration was performed without follicular flushing using a single lumen 17-, 19- or 20-gauge needle with vacuum pump suction (100-120 mm Hg) to harvest follicular contents into a 15 ml round bottom polystyrene centrifuge tube.

In all cases, rapid rotation of the aspiration needle around its long axis when the follicle had collapsed provided a curettage effect to assist the release of COCs. Although follicles were not flushed, the aspiration needle was removed from the subject and flushed frequently throughout the oocyte retrieval procedure to limit clotting and needle blockages.

Follicular aspirates were examined in the laboratory using a dissecting microscope. Aspirates tended to include more blood than typical IVF follicle aspirations, so samples and needles were washed with HEPES media (G-MOPS Plus, Vitrolife) to minimize clotting. Often, the aspirate was additionally filtered using a 70-micron cell strainer (Falcon, Corning) to improve the oocyte search process yield and speed. COCs were transferred using a sterile Pasteur pipette to a dish containing a pre-incubation LAG Medium (Medicult, Cooper Surgical) until use in the IVM procedure. The number of COCs aspirated was equal to roughly 40 to 50% of the subject’s AFC.

### Preparation of Ovarian Support Cells (OSCs)

OSCs were created from human induced pluripotent stem cells (hiPSCs) according to a 5-day transcription factor (TF)-directed protocol described previously.^41^ The cell lines utilized in this study came from a single original cell source consented for research use and were not generated on a per subject basis. The OSCs were produced in multiple batches and cryopreserved in vials of 120,000 to 150,000 live cells and stored in liquid nitrogen in CryoStor CS10 Cell Freezing Medium (StemCell Technologies). For the purposes of this study, the OSCs utilized in the OSC-IVM condition were generated through use of the transcription factors *NR5A1, RUNX2*, and *GATA4*. In Experiment 1, an additional control cell line of fetal ovarian somatic cells (FOSC-IVM) was generated using overexpression of the transcription factors *NR5A1, RUNX1, GATA4*, and *FOXL2*. All OSC lines were negative for mycoplasma and human-infectious pathogens, had a freeze-thaw viability greater than 80%, were confirmed for cell identity via flow cytometry, immunofluorescence and/or qPCR of FOXL2, CD82, and NR2F2, and validated as steroidogenic via estradiol (E2) and progesterone (P4) ELISA assays (R&D Systems).

Culture dishes (4+8 Well Dishes, BIRR) for oocyte maturation experiments were prepared with culture media and additional constituents in 100μl droplets under mineral oil the day before oocyte collection, and equilibrated in the incubator overnight. The morning of oocyte collection, cryopreserved OSCs were thawed for 2-3 minutes at 37°C (in a heated bead or water bath), resuspended in OSC-IVM medium and washed twice using centrifugation pelleting at 300 x g for 5 minutes to remove residual cryoprotectant. Equilibrated OSC-IVM media was used for final cell resuspension. OSCs were then plated at a concentration of 100,000 cells per 100μl droplet by replacing 50μl of the droplet with 50μl of the OSC suspension 2-4 hours before the addition of oocytes to allow for culture equilibration and media conditioning (Figure S1). All FOSCs utilized in Experiment 1 were prepared in the same manner as OSCs.

### *In vitro* maturation

COCs were maintained in preincubation LAG Medium (Medicult, Cooper Surgical) at 37°C for 2-3 hours after retrieval prior to introduction to *in vitro* maturation conditions.

Three experimental comparisons were performed to address the following goals:

#### Experiment 1 (OSC activity)

The purpose of this comparison was to determine whether the stimulated OSCs were the active ingredient of the co-culture system versus other medium components alone or other cell types. Media were prepared following the manufacturer’s recommendations (Medicult, Cooper Surgical), and further supplemented with androstenedione and doxycycline (both necessary for activation of OSCs) in order to compare oocyte maturation outcomes with or without OSCs in the same medium formulation (Table 1).

**Table 1:** Cell Culture Media Conditions.

|  | EXPERIMENT 1:<br>OSC activity |  | EXPERIMENT 2:<br>OSC clinical relevance<br>*IVM only* |  | EXPERIMENT 3:<br>OSC clinical relevance<br>*embryo formation* |  |
| --- | --- | --- | --- | --- | --- | --- |
|  | *Experimental Groups* | *Control Group* | *Experimental Group* | *Control Group* | *Experimental Group* | *Control Group* |
| IVM Medium (Medicult, Cooper Surgical) | X | X | X | X | X | X |
| 75 mIU/ml of recombinant FSH (Millipore, F4021) | X | X | X | X | X | X |
| 100 mIU/ml of recombinant hCG (Sigma, CG10) | X | X | X | X | X | X |
| 500 ng/ml Androstenedione (Sigma, A-075I) | X | X | X |  | X |  |
| 1.0 µg/ml Doxycycline (StemCell Tech., 100-1047) | X | X | X |  | X |  |
| 100,000 cells per 100 µl droplet | X |  | X |  | X |  |

#### Experiment 2 (OSC clinical relevance, IVM only)

The purpose of this experiment was to compare the efficacy of the OSC-IVM system and the commercially available *in vitro* maturation system (Medicult IVM). The Control condition was prepared and supplemented by following the manufacturer’s recommendations (Medicult, Cooper Surgical), while medium for OSC-IVM was prepared with all supplements (Table 1).

#### Experiment 3 (OSC clinical relevance, embryo formation)

The purpose of this experiment was to compare the efficacy of the OSC-IVM system and the commercially available *in vitro* maturation system (Medicult IVM) in maturing human oocytes and generating euploid blastocysts. The Control condition was prepared and supplemented by following the manufacturer’s recommendations (Medicult IVM, Cooper Surgical), while medium for OSC-IVM was prepared with all supplements (Table 1). Mature oocytes from both groups were then treated identically for subsequent embryo formation steps.

### Subject description

#### Experiment 1

We collected 179 oocytes from 27 subjects who underwent abbreviated gonadotropin stimulation, with 49 utilized in OSC-IVM co-culture, 47 utilized in FOSC-IVM co-culture and 83 utilized in control culture. Co-culture in the Experimental and Control Conditions was performed in parallel when possible leading to excess control samples compared to individual experimental conditions, and COCs were distributed equitably when performed in parallel, resulting in 41 unique oocyte culture setups. Equitable distribution means that COCs with distinctly large cumulus masses, small cumulus masses, or expanded cumulus masses were distributed equally between the conditions. The COCs were distributed randomly between one or two conditions, as the low number of oocytes retrieved per subject was too few to distribute oocytes between all three conditions. COCs were subjected to *in vitro* maturation at 37°C for 24-28 hours in a tri-gas incubator with CO_2_ adjusted so that the pH of the bicarbonate-buffered medium was 7.2-7.3 and with the O_2_ level maintained at 5%.

#### Experiment 2

23 subjects were recruited, yielding 143 COCs with 70 utilized in the commercial IVM control and 73 utilized in the OSC-IVM condition. 2 subjects were excluded from analysis due to low (<2) or no oocytes retrieved, preventing pairwise assessment. Co–culture in the Experimental and Control Conditions was performed in parallel for all subjects. COCs were distributed equitably between the two conditions, as described above. COCs were subjected to *in vitro* maturation at 37°C for 28 hours in a tri-gas incubator with CO_2_ adjusted so that the pH of the bicarbonate-buffered medium was 7.2-7.3 and with the O_2_ level maintained at 5%.

#### Experiment 3

*In vitro* maturation with subsequent embryo formation was performed to assess developmental competence of the oocytes treated in the OSC-co-culture system in comparison to oocytes treated with commercially available IVM medium. 17 oocyte donors and 3 sperm donors were utilized for IVM with subsequent ICSI and blastocyst formation. 3 donors were excluded from analysis due to insufficient eggs retrieved during stimulation (≤ 2 oocytes). 131 COCs were included in the comparison, with 64 utilized in Commercial IVM control and 67 utilized in the OSC-IVM condition. Co–culture in the Experimental and Control Conditions was performed in parallel. COCs were distributed equitably, as described above. COCs were subjected to *in vitro* maturation at 37°C for 28 hours in a tri-gas incubator with CO_2_ adjusted so that the pH of the bicarbonate-buffered medium was 7.2-7.3 and with the O_2_ level maintained at 5%. Embryo formation proceeded in parallel, with culture proceeding no longer than day 7 post-IVM.

### Assessment of oocyte *in vitro* maturation and morphology

Following the 24-28 hour *in vitro* maturation culture, COCs were stripped of surrounding cumulus and corona cells via hyaluronidase treatment, then oocytes were assessed for maturation state according to the following criteria:

**GV** - presence of a germinal vesicle, typically containing a single nucleolus within the oocyte.

**MI** - absence of a germinal vesicle within the oocyte and absence of a polar body in the perivitelline space between the oocyte and the zona pellucida.

**MII** - absence of a germinal vesicle within the oocyte and presence of a polar body in the perivitelline space between the oocyte and the zona pellucida.

For Experiments 1 and 2, oocytes were individually imaged using brightfield microscopy on the ECHO Revolve. The images were later scored by a single trained embryologist according to the Total Oocyte Score (TOS) grading system.^42^ Oocytes were given a score of -1, 0, 1 for each of the following criteria: morphology, cytoplasmic granularity, perivitelline space (PVS), zona pellucida (ZP) size, polar body (PB) size, and oocyte diameter. ZP and oocyte diameter were measured using ECHO Revolve Microscope software and image analysis software Fiji (2.9.0/1.53t). The sum of all categories gave each oocyte a total oocyte score (ranging from -6 to +6) with higher scores indicating better morphological quality. Oocytes were individually placed in 0.2ml tubes containing 5μl Dulbecco’s Phosphate Buffered Saline (DPBS), flash frozen in liquid nitrogen and stored at -80°C for future molecular analysis.

For Experiment 3, individual oocyte imaging and scoring was not performed in order to minimize handling and disruption of oocytes prior to fertilization and embryo culture.

### *In vitro* fertilization and embryo culture

For Experiment 3, COCs were cultured as outlined above for 28 hours then denuded, assessed for MII formation and imaged as a group. Individual oocytes were inseminated via intracytoplasmic sperm injection (ICSI) on day 1 post-retrieval,^43^ then cultured in an Embryo Culture Medium (Global Total, Cooper Surgical, Bedminster, NJ) at 37°C in a tri-gas incubator with CO_2_ adjusted so that the pH of the bicarbonate-buffered medium was 7.2-7.3 and with the O_2_ level at 5%. 16 to 18 hours post-ICSI, fertilization was assessed and zygotes were imaged, and fertilized oocytes were cultured until day 3. Cleaved embryos were imaged and underwent laser-assisted zona perforation and further cultured to develop until the blastocyst stage.^44,45^ Blastocysts were assessed on day 5, 6, and/or 7, imaged and scored according to the Gardner scale, and then underwent trophectoderm biopsy for preimplantation genetic testing for aneuploidy (PGT-A) if deemed to be of usable quality (i.e. greater than or equal to a 3CC rating).^46^ Oocytes that failed to mature, failed to fertilize, failed to cleave, or failed to generate blastocysts of biopsy quality were flash frozen and sent for aneuploid analysis via PGT-A. No oocytes from this study were utilized or banked for transfer, implantation, or reproductive purpose.

Trophectoderm biopsies were transferred to 0.2ml PCR tubes and sent to an external research laboratory (Juno Genetics, Basking Ridge, NJ) for comprehensive chromosomal analysis using a single nucleotide polymorphism (SNP) based next generation sequencing (NGS) of all 46 chromosomes (pre-implantation genetic testing for aneuploidy, PGT-A).^47,48^

Whole remaining blastocysts from the TE-biopsy PGT-A tested samples were harvested for epigenetic analysis through enzymatic methylation sequencing (EM-Seq).

### Blastocyst Epigenetic Analysis

Five euploid day 5 or 6 blastocysts from the OSC-IVM condition and eight donated day 5 of 6 euploid blastocysts from a COS reference control were flash frozen in 5μl dPBS in 0.2ml PCR tubes and transferred to an external research laboratory for methylation sequencing preparation (Azenta/Genewiz, NJ). Blastocysts were individually harvested for high molecular weight genomic DNA (gDNA) extraction using the PicoPure DNA extraction kit (ThermoFisher) with an RNA carrier spike-in for yield improvement. Isolated gDNA was then prepared for epigenetic sequencing using the Enzymatic Methylation Sequencing Kit (EM-Seq) (NEB).^49^ Individual embryos were sequenced at a depth of 100 million reads using a 2 × 150 bp Illumina kit on the Illumina NovaSeq instrument. Methylation analysis was performed as briefly described. Methylation alignments were constructed using a well structured methylation analysis pipeline previously described (https://nf-co.re/methylseq/2.4.0).^50^ Briefly, bismarck was utilized to map to the human reference genome (Hg37), selected to match to reference data.^51,52^ A list of germline differentially methylated regions (gDMRs) was obtained from previous studies,^51^ and percent 5-methylcytosine (5mC) was calculated for each gDMR and plotted versus external literature controls. Additionally, whole genome 5mC levels were calculated and compared to the COS internal reference blastocysts as a model of standard fertility treatment embryos.

### Data analysis and statistics

Oocyte maturation outcome data was analyzed using Python statistical packages pandas (1.5.2), scipy (1.9.3), and statsmodels (0.13.5). Maturation percentages and embryo formation outcomes by donor group were analyzed by *t-test* as functions of the IVM environment (OSC-IVM or Media control). Two-tailed *T-test* statistics were computed comparing experimental versus control with Welch’s correction for unequal variance paired by donor. When comparing three independent groups one way analysis of variance (ANOVA) was utilized. Embryo formation outcomes were analyzed by logistic regression, comparing outcomes per COC subjected to IVM (OSC-IVM vs Control), significance was based on the OSC-IVM distribution compared to the Control mean. Bar graphs depict mean values and error bars represent standard error of the mean (SEM). Number of independent oocytes for the experiment, and paired versus unpaired *t-test* parameters are indicated in the figure legends and Materials and Methods.

### Prospective Sample Size Estimation

Experiment 1 was a pilot analysis of the OSC-IVM and control culture conditions to determine the mean maturation rate and standard error. Based on these outcomes, we projected that 20 subjects and/or 80 oocytes per group would yield sufficient sample size for statistical measurement of oocyte maturation rate with an 80% power. Based on the outcomes of Experiment 1 and 2, for Experiment 3 we estimated that 20 donors and/or a minimum of 65 oocytes in each group would allow for sufficient sample size for statistical measurement of euploid embryo formation with an 80% power.

## Results

### hiPSC-derived OSCs effectively promote human oocyte maturation in co-culture system

To obtain immature COCs, we utilized similar protocols to previous IVM studies, truncated IVF or hCG primed-IVM: abbreviated gonadotropin stimulation (3-4 days) most often with an hCG trigger.^19,53,54^ This abbreviated stimulation program, particularly with higher doses of hCG (i.e. 10,000IU), yielded a mixed cohort of COCs with different magnitudes of cumulus expansion. In Experiment 1, the rate of expanded COCs utilized in IVM was 14% ± 5.7% SEM in OSC-IVM, 13% ± 5.9% SEM in FOSC-IVM, and 20% ± 7.2% SEM in Media Control IVM, which did not significantly differ between groups (*p*=0.7156, ANOVA). In Experiment 2, the rate of expanded COCs utilized in IVM was 12.3% ± 3.9% SEM in OSC-IVM and 12.3% ± 4.9% SEM in the Commercial IVM control, which did not significantly differ between groups (*p*=0.99, paired *t*-test). In Experiment 3, for both OSC-IVM and Commercial IVM control the rate of expanded COCs utilized in IVM was 3.17% ± 3.17%, which did not significantly differ between groups (*p*=0.99, paired *t*-test).

Oocyte donor demographics and treatment regimens are shown in Table 2 for each experimental group. Overall, the results demonstrate we were able to retrieve oocytes from donors, albeit at a lower yield than traditional controlled ovarian hyperstimulation cycles. Follicle measurements taken during stimulation in all subjects shows extraction occurred in follicles less than 6mm on average 33.03% ± 3.02% SEM, between 6 mm and 12 mm 64.87% ± 3.16% SEM, and above 12 mm 2.09% ± 0.63% SEM. Serum levels of E2, P4, and LH at OPU were similar in donors between groups within Experiment 1, but different across Experiments 1, 2 and 3. These hormonal values show evidence of early follicular growth with minimal instances of premature luteinization (Table 2).

**Table 2:** Donor Demographic and Stimulation Characteristics.

|  | **EXPERIMENT 1:**<br>**OSC activity** |  |  | **EXPERIMENT**<br>**2:**<br>**OSC clinical**<br>**relevance** | **EXPERIMENT**<br>**3:**<br>**OSC clinical**<br>**relevance** |
| --- | --- | --- | --- | --- | --- |
|  | *OSC-IVM*<br>*Group* | *FOSC-IVM*<br>*Group* | *Control*<br>*Group* | *IVM only* | *Embryo*<br>*Formation* |
| Number of oocyte donors included | 12 | 11 | 18 | 21 | 14 |
| Oocyte donor age (Avg ± SEM) | 26 ± 0.94 | 28 ± 0.83 | 24 ± 1.1 | 29 ± 0.63 | 31.1 ± 0.69 |
| Oocyte donor BMI (Avg ± SEM) | 22 ± 1.2 | 22 ± 0.51 | 22 ± 0.52 | 22 ± 0.58 | 25.3 ± 1.50 |
| Total rFSH used for stimulation (IU) | 325-600 | 325-600 | 325-600 | 600 | 5 days Clomid plus 150-300 IU or 600 IU |
| hCG trigger utilized (IU) | 10,000 | 10,000 | 10,000 | 10,000 | 0 IU or 2,500 IU |
| COCs obtained (Mean ± SEM) | 7.8 ± 0.98 | 8.5 ± 1.01 | 7.7 ± 1.03 | 7.0 ± 0.64 | 9.5 ± 1.40 |
| AMH (ng/ml) (Mean ± SEM) | 5.1 ± 0.65 | 5.1 ± 0.81 | 4.2 ± 0.53 | 4.1 ± 0.42 | 6.4 ± 0.86 |
| AFC (Mean ± SEM) | 17.7 ± 1.28 | 16.7 ± 0.98 | 15.1 ± 1.09 | 20.1 ± 0.12 | 27.9 ± 2.05 |
| E2 at OPU (ng/ml) (Mean ± SEM) | 330 ± 52.4 | 308 ± 61.7 | 159 ± 52.5 | 474 ± 59.7 | 387 ± 135.4 |
| P4 at OPU (ng/ml) (Mean ± SEM) | 0.69 ± 0.12 | 0.71 ± 0.10 | 0.98 ± 0.16 | 1.02 ± 0.12 | 1.1 ± 0.17 |
| LH at OPU (ng/ml) (Mean ± SEM) | 4.35 ± 0.99 | 4.48 ± 0.90 | 5.27 ± 1.10 | 5.01 ± 0.90 | 8.75 ± 3.3 |

In Experiment 1, oocytes from each donor were allocated to two simultaneous culture conditions (control IVM and FOSC-IVM or OSC-IVM) when possible. For Experiment 2 and 3, the control and OSC-IVM arms contained paired sibling oocytes from each donor. A depiction of the three experimental designs is shown (Figure 1A-C, Figure S1), including a representative image of the OSC co-culture (Figure 1D).

We have previously demonstrated that hiPSC-derived OSCs are predominantly composed of granulosa-like cells and can be generated through overexpression of different transcription factor combinations.^41^ In response to hormonal stimulation treatment *in vitro*, the OSCs produce extracellular matrix remodelers, growth factors and steroids necessary for paracrine interaction with oocytes and cumulus cells.^41^ To investigate whether hiPSC-derived OSCs are functionally capable of promoting human oocyte maturation *in vitro*, we established a co-culture system of these cells with freshly retrieved cumulus-oocytes complexes (COCs) and assessed oocyte maturation rates after 24-28 hours (see Materials and Methods, Experiment 1). We likewise tested a fetal version of OSCs (FOSC-IVM) to determine if the adult OSC-IVM variant was specific in promoting maturation or if other cell co-cultures could likewise promote maturation.

In this comparison, due to low numbers of retrieved oocytes per donor, we were unable to consistently split oocytes between all conditions simultaneously. Therefore each group contains oocytes from predominantly non-overlapping donor groups and pairwise comparisons are not utilized. Strikingly, we observed a statistically significant improvement (∼1.5x) in maturation outcomes for oocytes that underwent IVM with OSC-IVM (Figure 2A) compared to the Media Control. More specifically, the OSC-IVM group yielded a maturation rate of 68% ± 6.83% SEM versus 46% ± 8.51% SEM in the Media Control (Figure 2A, *p*=0.02592, unpaired *t-*test). The FOSC-IVM did not yield significant improvement in maturation rate over the Media Control, yielding 51% ± 9.23% SEM (Figure 2A, *p*=0.77 unpaired *t*-test). These results support the functional activity of specifically hiPSC-derived adult-like OSCs significantly improving oocyte maturation using this *in vitro* co-culture system. We next examined whether hiPSC-derived OSCs would also affect the outcome of oocyte morphological quality as assessed by the Total Oocyte Score (TOS). Interestingly, the assessment scores were not statistically significantly different for the three groups (Figure 2B; ANOVA, *p*=0.3728), indicating that the mature MII oocytes were of equivalent morphological quality between the IVM conditions. Altogether, these data indicate that OSC co-culture improves human oocyte maturation without a detrimental effect on morphological quality and highlights the potential for the use of hiPSC-derived OSCs as a high performing system for cumulus enclosed oocyte IVM.

**Figure 2:**
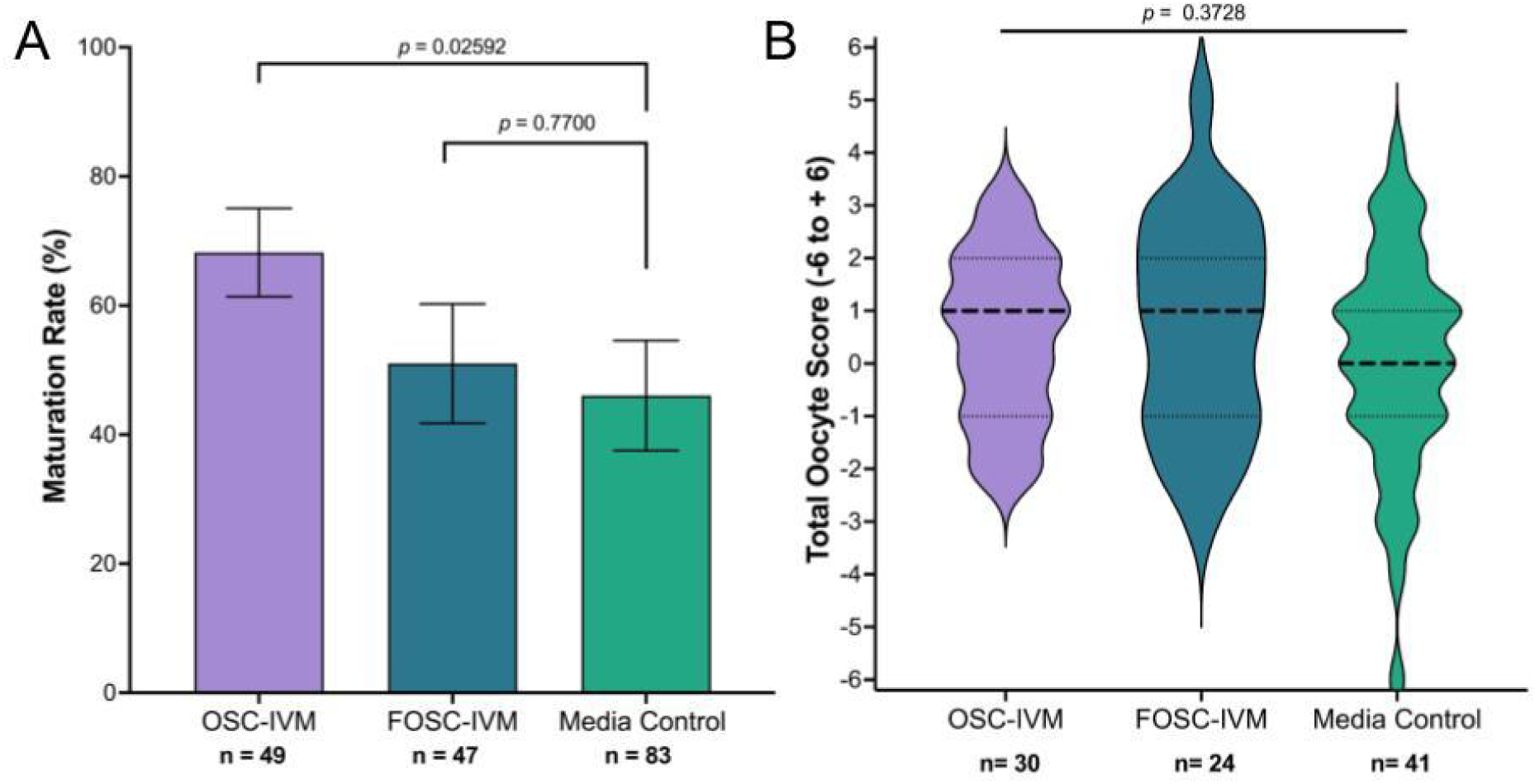
OSCs improve human oocyte maturation rates compared to supplemented IVM medium lacking OSCs or containing other cell types. **A** Maturation rate of oocytes after 24-28 hour IVM experiments in Experiment 1, including oocytes co-cultured with OSCs, FOSCs, or in Media Control. *n* indicates the number of individual oocytes in each culture condition. Error bars indicate mean ± SEM. *p*-value derived from unpaired *t-*test comparing OSC-IVM to control (Media Control) and FOSC-IVM to control (Media Control). **B** Total Oocyte Score (TOS) generated from imaging analysis of MII oocytes after 24-28 hour IVM experiments. *n* indicates the number of individual MII oocytes analyzed. Median (dashed line) and quartiles (dotted line) are indicated. A one way ANOVA indicated no significant difference between the means of the three groups. Due to low numbers of retrieved oocytes per donor, oocytes could not be consistently split between both conditions. Groups contain oocytes from predominantly non-overlapping donor cohorts thus pairwise comparisons are not utilized.

### Oocyte maturation rates in OSC-IVM outperforms commercially available IVM system

To further examine the potential for using OSC-IVM as a viable system to mature human oocytes in a more clinical setting, we compared our OSC co-culture system against a commercially available IVM standard, used according to the manufacturer’s instructions for use, with no modification (Medicult IVM, Cooper Surgical). We performed a sibling oocyte study comparing the MII formation rate and oocyte morphological quality after 28 hours of *in vitro* maturation (Materials and Methods, Experiment 2). Notably, OSC-IVM yielded a statistically significant ∼1.6x higher average MII formation rate (68% ± 6.74% of mature oocytes in OSC-IVM versus 43% ± 7.90% in the control condition; Figure 3A, *p*=0.0349, paired *t-*test). Similar to our prior observations, co-culture with hiPSC-derived OSCs did not affect oocyte morphological quality as measured by TOS, indicating equivalent oocyte morphological characteristics (Figure 3B, p=0.9420, unpaired *t-*test). These results show that OSC-IVM significantly outperformed the commercially available IVM culture medium in MII formation rate with no apparent detriment to oocyte morphological quality, pointing to a beneficial application of this product for human IVM.

**Figure 3:**
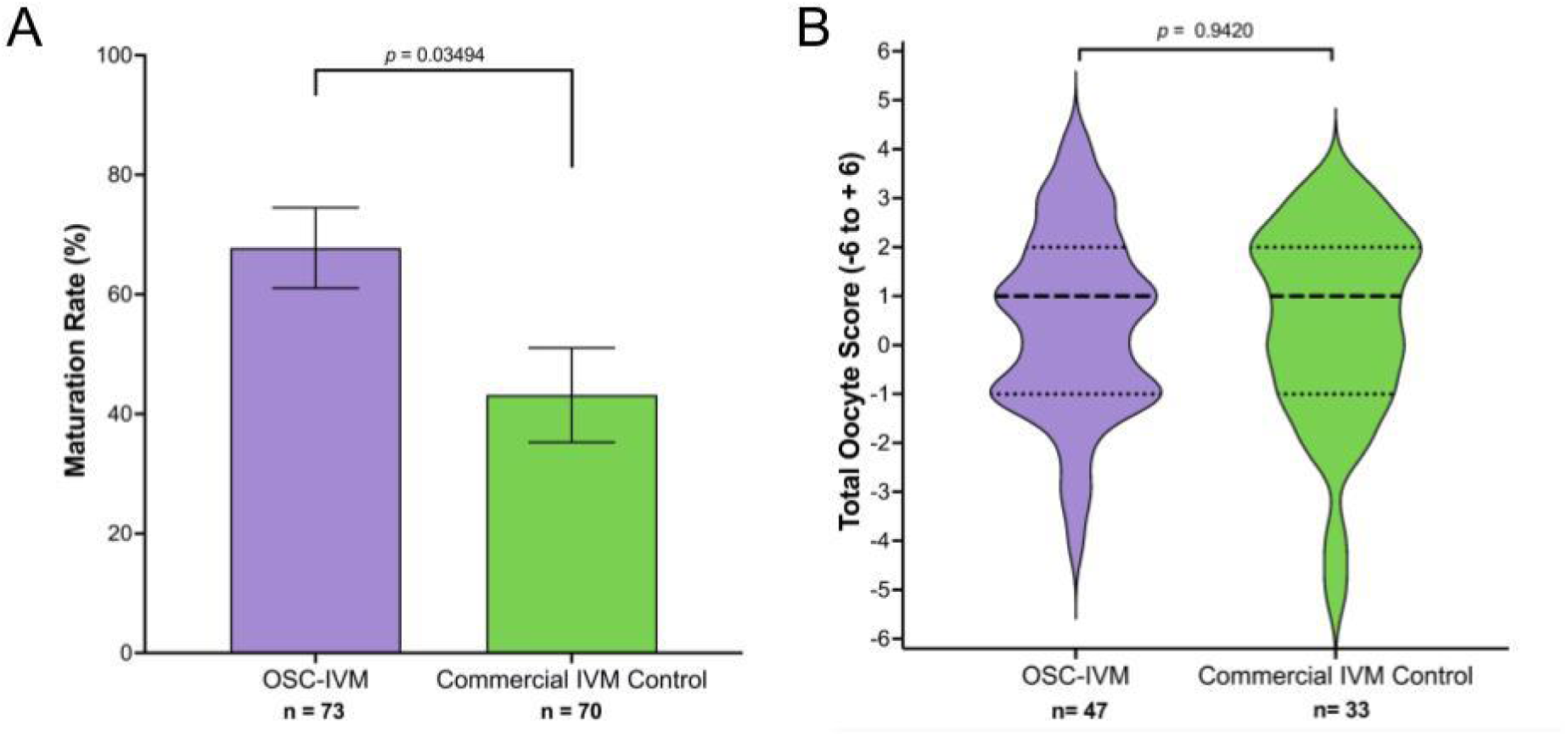
OSC-IVM demonstrates improved oocyte maturation compared to a commercially available IVM system. **A** Maturation rate of oocytes after 28 hour IVM experiments in Experiment 2, including oocyte co-culture with OSCs or in Commercial IVM Control. *n* indicates the number of individual oocytes in each culture condition. Error bars indicate mean ± SEM. p-value derived from paired *t-test* comparing Experimental OSC-IVM to Commercial IVM Control. **B** Total Oocyte Score (TOS) derived from imaging analysis of MII oocytes after 28 hour IVM experiments. *n* indicates the number of individual MII oocytes analyzed. Median (dashed line) and quartiles (dotted line) are indicated. An unpaired *t-test* indicated no significant (*p*=0.9420) difference between the means. COCs from each donor were randomly and equitably distributed between control and intervention to allow for pairwise statistical comparison.

### Cumulus enclosed immature oocytes from abbreviated gonadotropin stimulation matured by OSC-IVM are developmentally competent for embryo formation

We next investigated the developmental competency of oocytes treated in the OSC-IVM system by assessing euploid blastocyst formation after insemination, compared to the commercially available IVM control. Utilizing a limited cohort of 17 subjects who underwent abbreviated stimulation (see Materials and Methods Experiment 3, Table 1, and Table 2), we investigated whether OSC-IVM treated oocytes were capable of fertilization, cleavage, and formation of euploid blastocysts. Of the 17 subjects, 3 were excluded due to low egg retrieval yield (≤2 oocytes), which prevented pairwise comparison. As expected, OSC-IVM yielded a significantly higher ∼1.3x higher average MII formation rate with 56% ± 9.2% of mature oocytes compared to 43% ± 6.4% in the control condition (*p*=0.0476, logistic regression) (Table 3). Matured oocytes from both conditions were immediately utilized for ICSI and followed until embryo formation (blastocyst stage). Per MII oocyte, OSC-IVM oocytes display similar fertilization and cleavage rates (74% ± 10.2% and 74% ± 10.2%) versus the control (81% ± 9.3% and 73% ± 11.3%). OSC-IVM MIIs trended towards improved blastocyst formation (64% ± 10.2%) versus control (50.7% ± 11.3%). OSC-IVM yielded a trend towards improvement in day 5 or 6 blastocyst formation per COC (28% ± 7.1%) compared to the commercial IVM control (15% ± 4.4%) (Figure 4, Table 3). Strikingly, OSC-IVM yielded a statistically significant improvement in day 5 or 6 euploid blastocyst formation per COC (25% ± 7.5%) compared to the commercial IVM control (11% ± 3.7%) (Figure 4, Table 3). These findings demonstrate that OSC-IVM generates healthy matured oocytes with high quality developmental competency. These results additionally demonstrate OSC-IVM is capable of producing healthy, euploid embryos from abbreviated stimulation cycles at a higher rate than the commercially available IVM condition, highlighting the clinical relevance of this novel system for IVM ART practice.

**Table 3:**
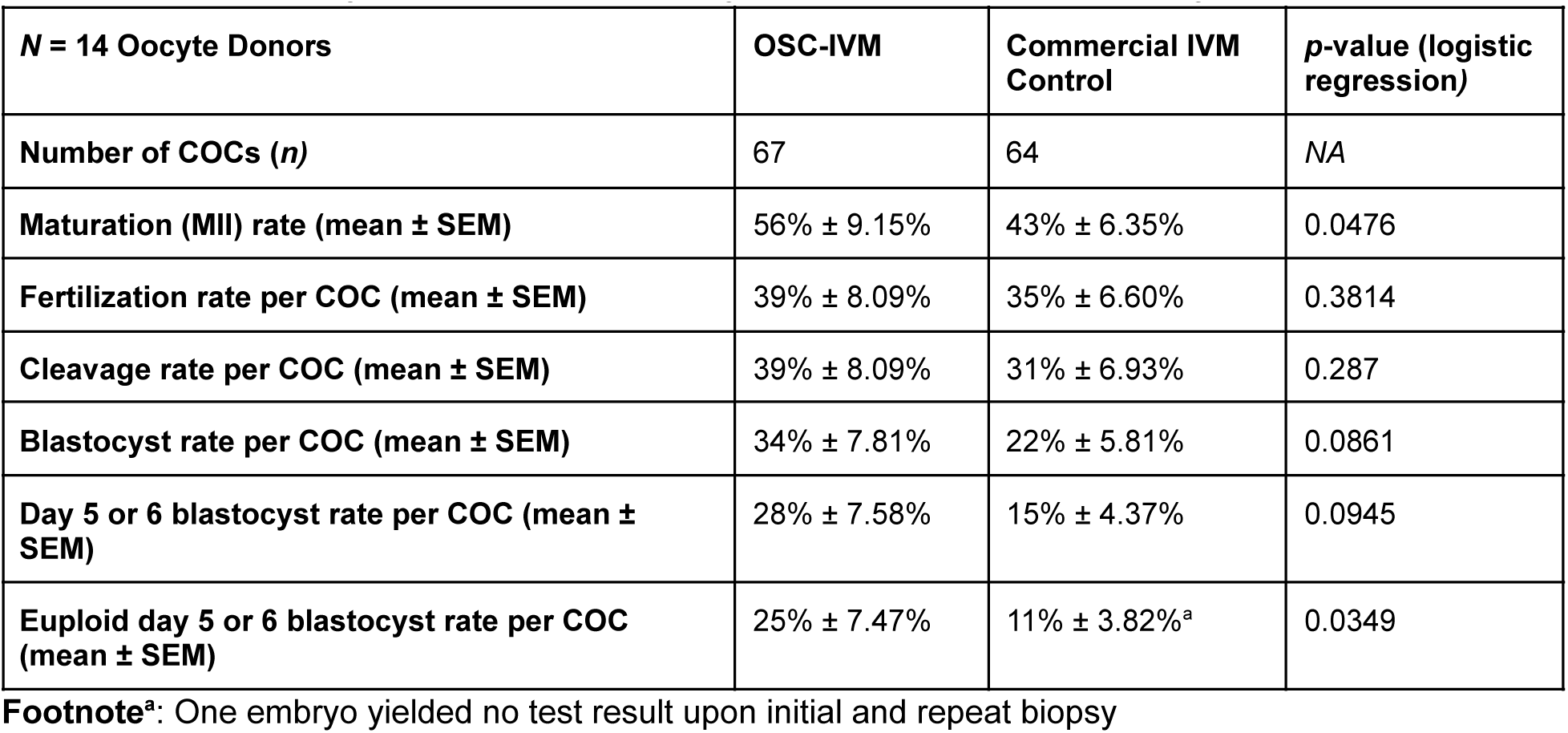
OSC-IVM oocytes are developmentally competent for euploid embryo formation.

**Figure 4:**
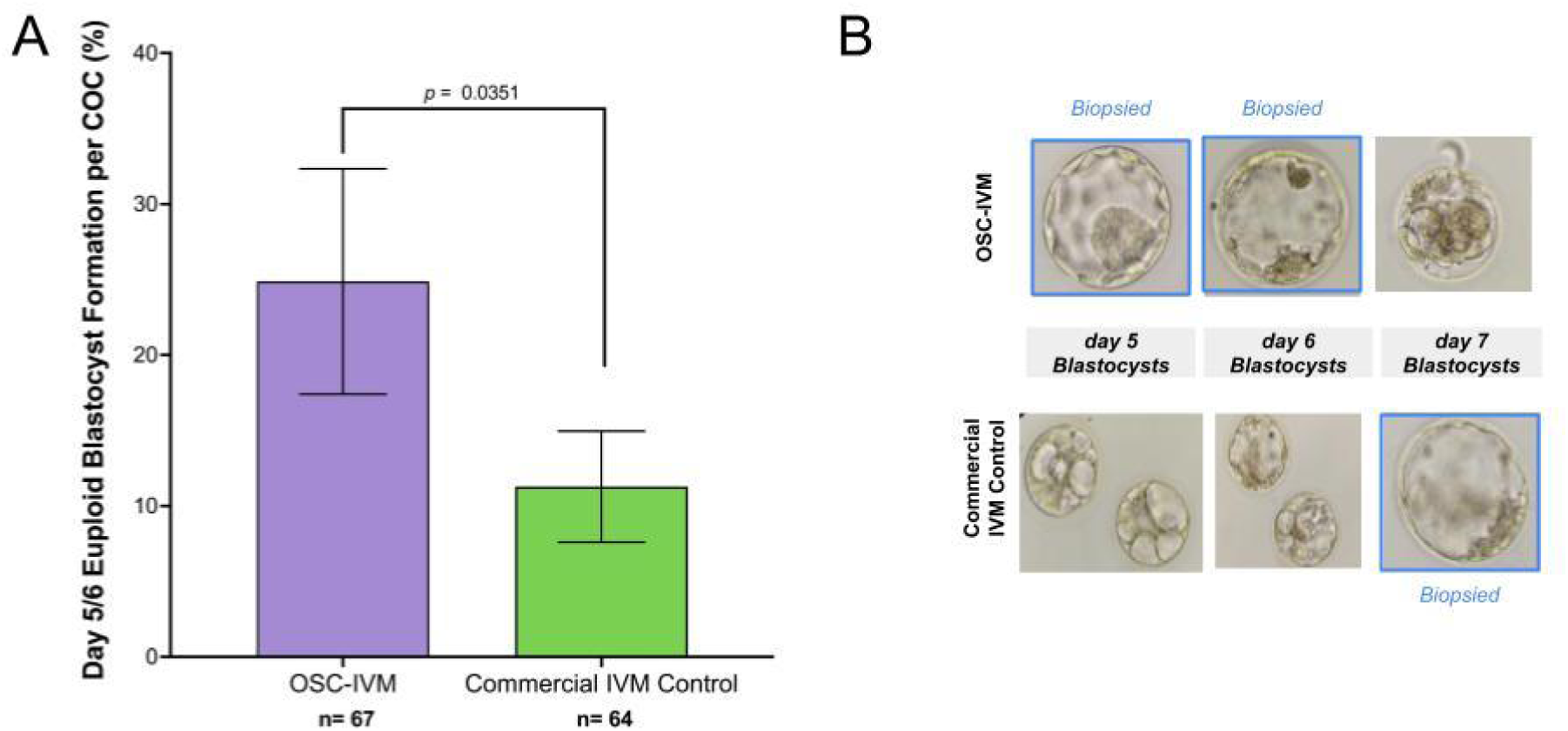
OSC-IVM assisted oocytes are developmentally competent for healthy embryo formation. **A** Euploid blastocyst (day 5 or 6) formation rate per COC in Experiment 3 after oocyte co-culture with OSCs or in Commercial IVM Control. *n* indicates the number of individual oocytes in each culture condition. Error bars indicate mean ± SEM. p-value derived from logistic regression comparing Experimental OSC-IVM to control (Commercial IVM Control). **B** Representative images of embryo formation in OSC-IVM versus Commercial IVM conditions at day 5, 6, and 7 of blastocyst formation. Embryos that were of suitable vitrification quality are labeled as “biopsied” and were utilized for trophectoderm biopsy and PGT-A.

### Euploid blastocysts from OSC-IVM show normal epigenetic profile compared to COS and CAPA-IVM

Using five euploid day 5 or 6 blastocysts generated from OSC-IVM and an additional eight donated controls obtained from conventional controlled ovarian stimulation (COS) and treatment, we assessed embryo epigenetic quality through enzymatic methylation sequencing (EM-Seq). We further utilized external reference data generated on COS and other IVM approaches (CAPA-IVM) as further comparison controls.^51^ Our results show OSC-IVM generates embryos with no significant difference in 5 methylcytosine (5mC) methylation profile across 19 well characterized germline differentially methylated regions (gDMRs) (Figure 5A, ANOVA with multiple comparisons tests, *p*=0.2807). Furthermore, whole genome 5mC profiling shows that OSC-IVM embryos show no significant difference compared to COS internal reference controls (Figure 5B, unpaired t-test, *p*=0.7970). Together, these results indicate that OSC-IVM is not only capable of improving oocyte maturation and improving euploid day 5 and 6 blastocyst formation, but it also generates embryos with global epigenetic profiles similar to embryos generated from conventional COS and other IVM treatments.

**Figure 5:**
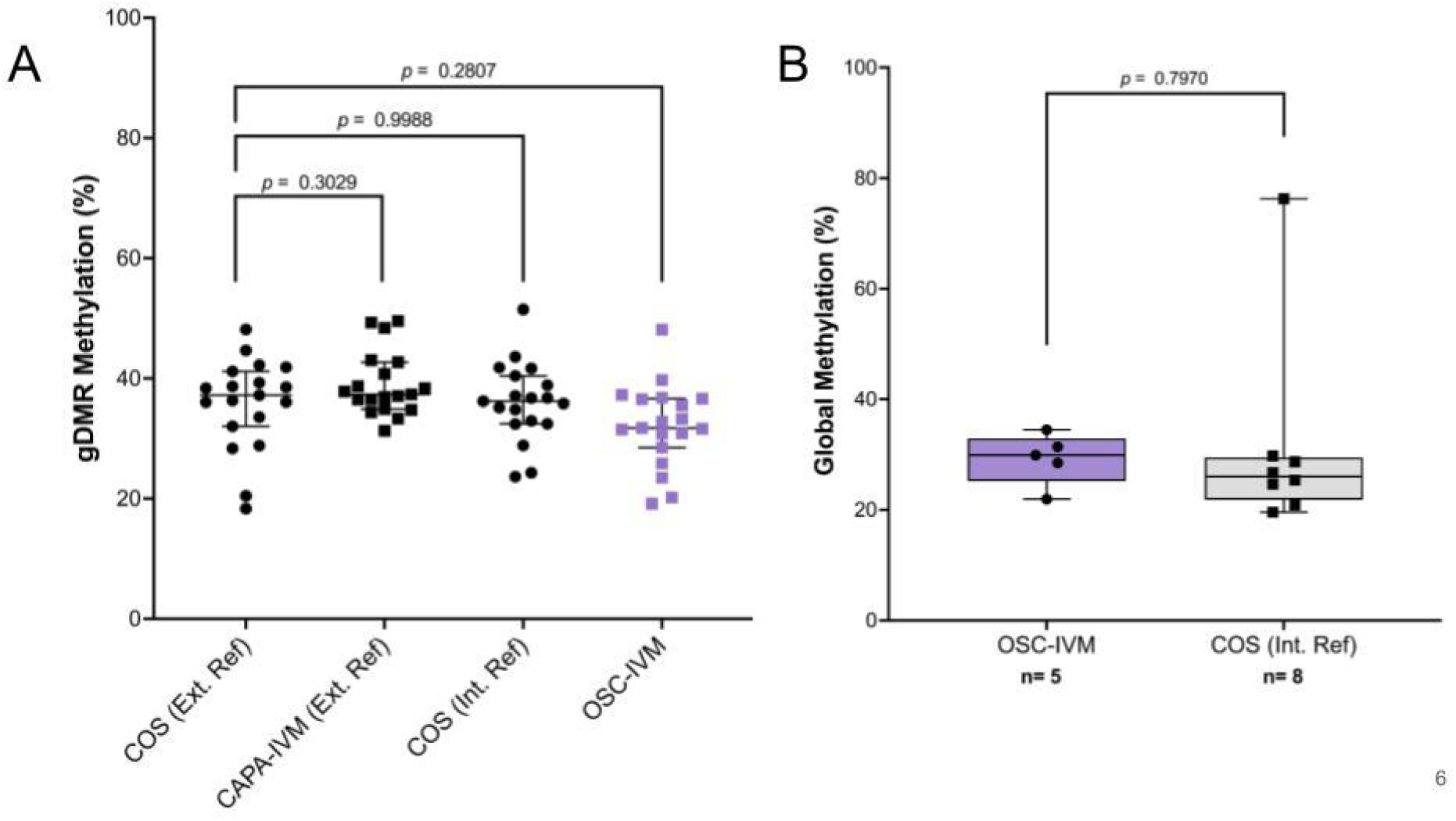
OSC-IVM blastocysts display similar global and germline methylation profiles to COS blastocysts. **A** 19 germline differentially methylated regions (gDMRs) were profiled for their methylation pattern in five OSC-IVM day 5 or 6 euploid blastocysts and eight donated day 5 or 6 euploid blastocysts after conventional stimulation (COS). Additional control data was obtained from Saenz-de-Juano et al. 2019 for CAPA-IVM and COS samples. Symbols represent gDMR percentages of individual embryos. Data is plotted as median with error bars representing 95% CI. Statistical significance testing was performed using one way ANOVA with multiple comparisons testing for post hoc analysis comparison to the external reference COS control. **B** Whole genome methylation analysis of 5-methylcytosine (5mC) levels was analyzed for the five OSC-IVM blastocysts and eight donated COS blastocyst controls. Statistical significance testing was performed using two-way unpaired t-test comparison to the external reference COS control.

## Discussion

It is well established that immature oocytes obtained from human ovarian follicles can mature upon extraction and *in vitro* culture.^27,55^ However, the rate and extent of oocyte maturation using these conditions are not optimized, resulting in highly unsynchronized nuclear and cytoplasmic maturation, and leaving IVM as an underperforming and underutilized technology. Current commercial IVM media have incrementally improved this process. With further optimization and development of additional culture tools, it is possible to improve the rate and extent of oocyte maturation, yielding high quality oocytes capable of healthy embryo formation. In this study, we investigate a novel IVM approach using human stem cell-derived ovarian support cells as a co-culture system for IVM of human oocytes. We demonstrated that this OSC-IVM system outperforms media without OSCs and other cell lines as well as a commercially available IVM culture media for MII formation. This work is the first of its kind to explore the potential of stem cell-derived OSCs as a tool for IVM of human COCs and shows the value of this and other cell engineering approaches for application in fertility treatment. This study also demonstrated that the OSCs specifically, and not just cell co-culture in general, provided a benefit to the oocytes beyond that of the acellular supplements to the culture conditions (hormones, steroids, doxycycline). Therefore, the engineered granulosa-like cells are a beneficial addition to the *in vitro* culture environment for oocytes.

Notably, many previous IVM studies stopped at the embryo cleavage stage, whereas present ART practice increasingly relies heavily upon culture to the blastocyst stage and PGT-A as an embryo selection tool. Our study employs these more advanced selection tools. The OSC-IVM system results in MII oocytes with a high degree of developmental competence, as demonstrated by their ability to achieve fertilization, become embryos capable of cell divisions, and form blastocysts characterized with a normal euploid complement of chromosomes and normal epigenetic profile. In short, oocytes matured in the presence of OSCs are capable of becoming healthy euploid embryos that would likely be competent to implant and become live born babies. The ability to improve the formation of healthy, euploid blastocysts is further evidence of the ability of the OSC-IVM technique to promote cytoplasmic maturation as well as nuclear maturation of oocytes, a key requirement for clinical utility of IVM systems. Further studies will explore this hypothesis. The high euploidy rate is a key indication of the potential clinical utility of OSC-IVM compared to conventional IVM systems.

This method of IVM involving co-culture integrates readily with existing procedures employed in ART laboratories, requiring no special equipment and only minimal training for clinical embryologists. The OSC-IVM approach is generated from a single source of highly qualified, commercially available hiPSCs and does not require patient-specific generation. This approach would therefore be highly scalable, reproducible and cost-effective as a novel IVM treatment for patients as opposed to individualized primary cell co-culture or conventional COS treatment. Likewise, OSC-IVM performs effectively in hCG triggered cycles with variable follicle size, which is an improvement over other IVM systems that are ill-suited for these more common stimulation styles.^19^ Our results show that OSC-IVM can be utilized across a variety of stimulation regimens, follicle extraction methodologies, and without hCG triggers, making the system highly amenable to customized abbreviated stimulation programs across the spectrum from strict IVM to truncated IVF practice. Additionally, the paracrine interaction between the COCs and OSCs in this system, in which direct connections/gap junctions are not formed, affords a high degree of flexibility in co-culture format delivery and allows for a broad range of potential culture configurations.

This study represents an important first step in establishing stem cell derived OSCs as a co-culture platform for IVM. This study builds on previous findings that demonstrate a benefit to using primary granulosa cell co-culture for *in vitro* maturation of oocytes.^37–40^ The use of OSCs derived from hiPSCs represents a powerful approach for providing curated granulosa-like cells with characteristics more similar to a follicular rather than the luteal phase environment, as well as providing a highly controlled and robust manner for their consistent production and qualification for broad use in clinical settings. Indeed, recent advances in IVM practice such as CAPA-IVM, which seeks to mimic aspects of granulosa cell function in the IVM culture environment through supplementation of CNP and AREG show that the biological role of granulosa cells in maturing oocyte *in vitro* is paramount to improving oocyte maturation rate and oocyte quality.^11,19^ Those findings are largely consistent with the findings of this study that show direct supplementation of granulosa-like cells indeed improves oocyte maturation and developmental competence. While a direct comparison of OSC-IVM and CAPA-IVM was not performed in this study, the rates of oocyte maturation were similar for both OSC-IVM and CAPA-IVM as well as for the common commercial IVM control used in both studies. OSC-IVM results in a significant improvement in day 5 and 6 euploid blastocyst formation rate, which was not demonstrated in previous CAPA-IVM studies, which stopped at the cleavage stage of embryo development. Future studies are warranted to explore differences in maturation and embryo formation rates between OSC-IVM and other IVM approaches.

### Limitations and Reasons for Caution

In general, while this study establishes a novel co-culture system for improving IVM outcomes compared to currently available IVM methods, a number of limitations and areas of future research remain. The nature of this study as a basic research study limits the generalizability of the findings, and future randomized control clinical evaluations will be important for establishing the value of the approach. Given that COCs were utilized from hCG triggered cycles, the underlying maturation state of the oocytes prior to IVM could not be empirically assessed while maintaining the integrity of the cumulus-oocyte connections. Further, while equitable distribution of COCs between conditions was performed, slight imbalances in the GV/MI/MII input distribution may represent a limitation of the study. This study utilized relatively small sample sizes overall and larger studies of the approach will assist in clarifying the broader efficacy and safety. Further, additional studies should be performed to more precisely determine the mechanism of the approach and characterize the specific secreted factors and changes to the *in vitro* growth environment that are responsible for the improved IVM performance. While it is well established that granulosa cell co-culture improves oocyte maturation in COCs and denuded oocytes, it will still be beneficial to determine how these hiPSC-derived OSCs perform their function and compare them to primary cell co-culture systems.^37–39^ Lastly, important validation of the approach’s clinical utility will be the ability to generate healthy live births, and further clinical evaluation is warranted to determine if such approach provides clinical value for patients over existing IVM and conventional COS treatments.

## Acknowledgments

This work was performed with the support of clinical partnerships at Spring Fertility New York, Extend Fertility, Ruber Clinic of Madrid, Pranor Clinic of Lima, and ReproART clinic of Georgia. We thank the dedicated support and work of the embryology and support staff teams at these clinics for coordinating and managing the collaborative study. We additionally thank the Wyss Institute for Biologically Inspired Engineering at Harvard University for material transfer of reagents used in this preclinical work. We thank Professor Mary Herbert, Professor Phillip Jordan, Professor George Church, Professor Kristin Baldwin, Dr. Sara Vaughn, and Professor David Albertini for advice and guidance on the use of OSCs in IVM work. We also thank the New York University flow cytometry and imaging cores, Azenta/Genewiz, and Juno Genetics for their assistance in data generation and analysis.

## Author Contributions

C.C.K. designed, supervised, and coordinated the study as well as provided technical assistance in cell engineering, production, qualification, and sequencing. S.P, A.G., M.M., F.B., C.A., M.S., S.M., D.A.K., M.F., and M.F. performed all embryology work for the study. B.P., K.S.P., and A.N. produced and qualified OSC batches. G.R. performed all statistical analysis and embryo epigenetic analysis. A.F. and T.S. coordinated study logistics. D.H.M., K.W., N.D.T., and L.G., assisted in study design, stimulation concepts, and embryology workflows. M.P.S., P.R.J.F., and P.C. assisted in OSC data interpretation S.O., D.O., J.K. and P.K. performed all oocyte donor stimulations and retrievals. C.C.K., B.P., K.S.P., G.R., and D.H.M. wrote the manuscript with significant input from all authors.

## Supplemental Figures

**Figure S1:**
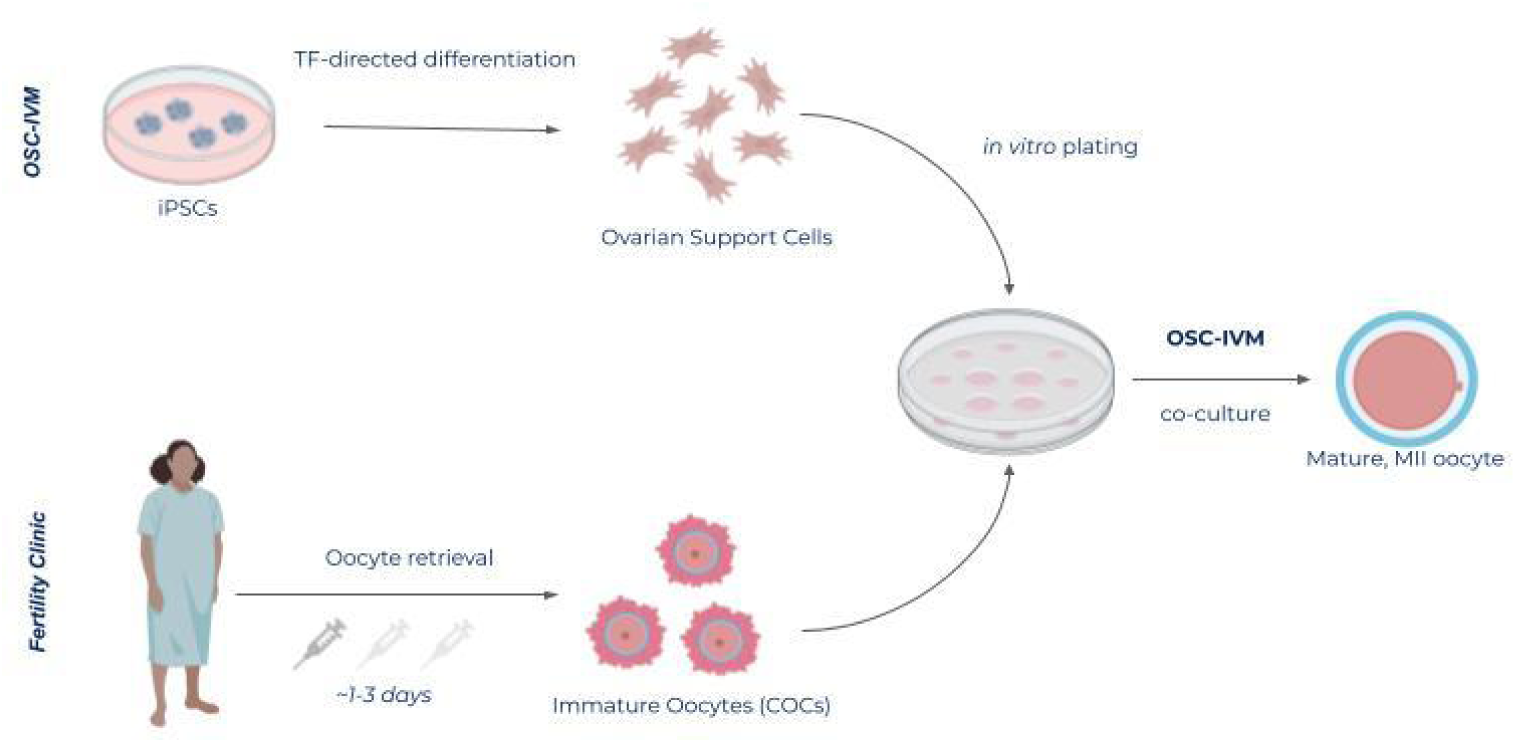
Ovarian support cells (OSC) co-culture system. Schematic of the experimental co-culture IVM approach. hiPSCs were differentiated using inducible transcription factor overexpression to form ovarian support cells (OSCs). Immature human cumulus oocyte complexes (COCs) were obtained from donors in the clinic after undergoing abbreviated gonadotropin stimulation. In the lab, embryology dishes were prepared including OSCs seeding as required, and COCs were introduced for IVM co-culture. Oocyte maturation and morphological quality were assessed after 24-28 hours IVM co-culture, and samples were either banked for analysis or utilized for embryo formation.

## Notes

**Funding Statement:** This work was funded by the for-profit entity Gameto Inc. and no other grant or funding agency.

**Capsule:** Induced pluripotent stem cell-derived ovarian support cell co-culture improves *in vitro* maturation of human oocytes obtained from abbreviated gonadotropin stimulation cycles, yielding healthy euploid blastocysts.

### Competing Interest Statement

The majority of this work was performed by employees and shareholders of the for-profit biotechnology company Gameto Inc. C.C.K., S.P., M.M., A.G., B.P., K.S.P., G.R., A.D.N., are listed on a patent covering the use of OSCs for in vitro maturation, Non-provisional Patent Application No. 63/492,210. Additionally, C.C.K. and K.W. are listed on two patents covering the use of OSCs for in vitro maturation: Non-provisional Patent Application No. 17/846,845 and U.S. Non-provisional Patent Application No. 17/846,725. C.C.K., M.P.S. and P.C. additionally are listed on a patent for the TF-directed production of granulosa-like cells from stem cells: U.S. Provisional Application No. 63/326,640.

### Summary of Updates

Update to Table 3 p-values and addition of an author who was not added to the revised manuscript in error.

## References

[1] NSFG - listing I - key Statistics from the National Survey of family growth. Published November 6, 2019. https://www.cdc.gov/nchs/nsfg/key_statistics/i.htm. Accessed June 1, 2022

[2] Gaskins AJ, Zhang Y, Chang J, Kissin DM. Predicted probabilities of live birth following assisted reproductive technology using United States national surveillance data from 2016 to 2018. Am J Obstet Gynecol. Published online January 23, 2023. doi:10.1016/j.ajog.2023.01.014

[3] Fauser BC. Towards the global coverage of a unified registry of IVF outcomes. Reprod Biomed Online. 2019;38(2):133–137.

[4] El Tokhy O, Kopeika J, El-Toukhy T. An update on the prevention of ovarian hyperstimulation syndrome. Womens Health. 2016;12(5):496–503.

[5] Ethics Committee of the American Society for Reproductive Medicine. Electronic address. Disparities in access to effective treatment for infertility in the United States: an Ethics Committee opinion. Fertil Steril. 2021;116(1):54–63.

[6] Braam SC, de Bruin JP, Mol BWJ, van Wely M. The perspective of women with an increased risk of OHSS regarding the safety and burden of IVF: a discrete choice experiment. Hum Reprod Open. 2020;2020(2):hoz034.

[7] De Vos M, Smitz J, Woodruff TK. Fertility preservation in women with cancer. Lancet. 2014;384(9950):1302–1310.

[8] De Vos M, Grynberg M, Ho TM, Yuan Y, Albertini DF, Gilchrist RB. Perspectives on the development and future of oocyte IVM in clinical practice. J Assist Reprod Genet. 2021;38(6):1265–1280.

[9] Walls ML, Hunter T, Ryan JP, Keelan JA, Nathan E, Hart RJ. In vitro maturation as an alternative to standard in vitro fertilization for patients diagnosed with polycystic ovaries: a comparative analysis of fresh, frozen and cumulative cycle outcomes. Hum Reprod. 2015;30(1):88–96.

[10] Zhang JJ, Merhi Z, Yang M, Bodri D, Chavez-Badiola A, Repping S, et al. Minimal stimulation IVF vs conventional IVF: a randomized controlled trial. Am J Obstet Gynecol. 2016;214(1):96.e1-e8.

[11] Vuong LN, Ho VNA, Ho TM, Dang VQ, Phung TH, Giang NH, et al. In-vitro maturation of oocytes versus conventional IVF in women with infertility and a high antral follicle count: a randomized non-inferiority controlled trial. Hum Reprod. 2020;35(11):2537–2547.

[12] Son WY, Tan SL. Laboratory and embryological aspects of hCG-primed in vitro maturation cycles for patients with polycystic ovaries. Hum Reprod Update. 2010;16(6):675–689.

[13] Sánchez F, Lolicato F, Romero S, De Vos M, Van Ranst H, Verheyen G, et al. An improved IVM method for cumulus-oocyte complexes from small follicles in polycystic ovary syndrome patients enhances oocyte competence and embryo yield. Hum Reprod. 2017;32(10):2056–2068.

[14] Grynberg M, Sermondade N, Sellami I, Benoit A, Mayeur A, Sonigo C. In vitro maturation of oocytes for fertility preservation: a comprehensive review. F&S Reviews. 2022;3(4):211–226.

[15] Mohsenzadeh M, Khalili MA, Anbari F, Vatanparast M. High efficiency of homemade culture medium supplemented with GDF9-β in human oocytes for rescue in vitro maturation. Clin Exp Reprod Med. 2022;49(2):149–158.

[16] De Vos M, Ortega-Hrepich C, Albuz FK, Guzman L, Polyzos NP, Smitz J, et al. Clinical outcome of non-hCG-primed oocyte in vitro maturation treatment in patients with polycystic ovaries and polycystic ovary syndrome. Fertil Steril. 2011;96(4):860–864.

[17] Guzman L, Ortega-Hrepich C, Albuz FK, Verheyen G, Devroey P, Smitz J, et al. Developmental capacity of in vitro-matured human oocytes retrieved from polycystic ovary syndrome ovaries containing no follicles larger than 6 mm. Fertil Steril. 2012;98(2):503–507.e1-e2.

[18] Vuong LN, L. AH, Ho VNA, Pham TD, Sanchez F, Romero S, et al. Live births after oocyte in vitro maturation with a prematuration step in women with polycystic ovary syndrome. J Assist Reprod Genet. 2020;37(2):347–357.

[19] Sanchez F, Le AH, Ho VNA, Romero S, Van Ranst H, De Vos M, et al. Biphasic in vitro maturation (CAPA-IVM) specifically improves the developmental capacity of oocytes from small antral follicles. J Assist Reprod Genet. 2019;36(10):2135–2144.

[20] Akin N, Le AH, Ha UDT, Romero S, Sanchez F, Pham TD, et al. Positive effects of amphiregulin on human oocyte maturation and its molecular drivers in patients with polycystic ovary syndrome. Hum Reprod. 2021;37(1):30–43.

[21] Braam SC, Consten D, Smeenk JMJ, Cohlen BJ, Curfs MHJM, Hamilton CJCM, et al. In Vitro Maturation of Oocytes in Women at Risk of Ovarian Hyperstimulation Syndrome-A Prospective Multicenter Cohort Study. Int J Fertil Steril. 2019;13(1):38–44.

[22] Shu-Chi M, Jiann-Loung H, Yu-Hung L, Tseng-Chen S, Ming-I L, Tsu-Fuh Y. Growth and development of children conceived by in-vitro maturation of human oocytes. Early Hum Dev. 2006;82(10):677–682.

[23] Pongsuthirak P, Songveeratham S, Vutyavanich T. Comparison of blastocyst and Sage media for in vitro maturation of human immature oocytes. Reprod Sci. 2015;22(3):343–346.

[24] Moschini RM, Chuang L, Poleshchuk F, Slifkin RE, Copperman AB, Barritt J. Commercially available enhanced in vitro maturation medium does not improve maturation of germinal vesicle and metaphase I oocytes in standard in vitro fertilization cases. Fertil Steril. 2011;95(8):2645–2647.

[25] Ma L, Cai L, Hu M, Wang J, Xie J, Xing Y, et al. Coenzyme Q10 supplementation of human oocyte in vitro maturation reduces postmeiotic aneuploidies. Fertil Steril. 2020;114(2):331–337.

[26] Fadini R, Dal Canto MB, Mignini Renzini M, Brambillasca F, Comi R, Fumagalli D, et al. Effect of different gonadotrophin priming on IVM of oocytes from women with normal ovaries: a prospective randomized study. Reprod Biomed Online. 2009;19(3):343–351.

[27] Edwards RG. Maturation in vitro of Mouse, Sheep, Cow, Pig, Rhesus Monkey and Human Ovarian Oocytes. Nature. 1965;208(5008):349–351.

[28] Straczynska P, Papis K, Morawiec E, Czerwinski M, Gajewski Z, Olejek A, et al. Signaling mechanisms and their regulation during in vivo or in vitro maturation of mammalian oocytes. Reprod Biol Endocrinol. 2022;20(1):37.

[29] The Ovary. Elsevier; 2019.

[30] Liu W, Xin Q, Wang X, Wang S, Wang H, Zhang W, et al. Estrogen receptors in granulosa cells govern meiotic resumption of pre-ovulatory oocytes in mammals. Cell Death Dis. 2017;8(3):e2662–e2662.

[31] Zhang H, Lu S, Xu R, Tang Y, Liu J, Li C, et al. Mechanisms of Estradiol-induced EGF-like Factor Expression and Oocyte Maturation via G Protein-coupled Estrogen Receptor. Endocrinology. 2020;161(12):bqaa190.

[32] Park JY, Su YQ, Ariga M, Law E, Jin SLC, Conti M. EGF-Like Growth Factors As Mediators of LH Action in the Ovulatory Follicle. Science. 2004;303(5658):682–684.

[33] Nilsson E, Skinner MK. Cellular interactions that control primordial follicle development and folliculogenesis. J Soc Gynecol Investig. 2001;8(1 Suppl Proceedings):S17–S20.

[34] Hsieh M, Zamah AM, Conti M. Epidermal growth factor-like growth factors in the follicular fluid: role in oocyte development and maturation. Semin Reprod Med. 2009;27(1):52–61.

[35] Fontana J, Martínková S, Petr J, Žalmanová T, Trnka J. Metabolic cooperation in the ovarian follicle. Physiol Res. 2020;69(1):33–48.

[36] Erickson GF, Shimasaki S. The physiology of folliculogenesis: the role of novel growth factors. Fertil Steril. 2001;76(5):943–949.

[37] Torre ML, Munari E, Albani E, Levi-Setti PE, Villani S, Faustini M, et al. In vitro maturation of human oocytes in a follicle-mimicking three-dimensional coculture. Fertil Steril. 2006;86(3):572–576.

[38] Johnson JE, Higdon HL 3rd, Boone WR. Effect of human granulosa cell co-culture using standard culture media on the maturation and fertilization potential of immature human oocytes. Fertil Steril. 2008;90(5):1674–1679.

[39] Jahromi BN, Mosallanezhad Z, Matloob N, Davari M, Ghobadifar MA. The potential role of granulosa cells in the maturation rate of immature human oocytes and embryo development: A co-culture study. Clin Exp Reprod Med. 2015;42(3):111–117.

[40] Zgórecka W, Jeseta M, Prochazka R, Amorim CA, Krajnik K, Mozdziak P, et al. Approaches for in vitro culture of granulosa cells and ovarian follicles. Medical Journal of Cell Biology. 2022;10(1):34–42.

[41] Pierson Smela MD, Kramme CC, Fortuna PRJ, Adams JL, Su R, Dong E, et al. Directed differentiation of human iPSCs to functional ovarian granulosa-like cells via transcription factor overexpression. Elife. 2023;12:e83291.

[42] Lazzaroni-Tealdi E, Barad DH, Albertini DF, Yu Y, Kushnir VA, Russell H, et al. Oocyte Scoring Enhances Embryo-Scoring in Predicting Pregnancy Chances with IVF Where It Counts Most. PLoS One. 2015;10(12):e0143632.

[43] Palermo GD, Neri QV, Schlegel PN, Rosenwaks Z. Intracytoplasmic sperm injection (ICSI) in extreme cases of male infertility. PLoS One. 2014;9(12):e113671.

[44] de Boer KA, Catt JW, Jansen RPS, Leigh D, McArthur S. Moving to blastocyst biopsy for preimplantation genetic diagnosis and single embryo transfer at Sydney IVF. Fertil Steril. 2004;82(2):295–298.

[45] McArthur SJ, Leigh D, Marshall JT, de Boer KA, Jansen RPS. Pregnancies and live births after trophectoderm biopsy and preimplantation genetic testing of human blastocysts. Fertil Steril. 2005;84(6):1628–1636.

[46] Gardner DK, Lane M, Stevens J, Schlenker T, Schoolcraft WB. Blastocyst score affects implantation and pregnancy outcome: towards a single blastocyst transfer. Fertil Steril. 2000;73(6):1155–1158.

[47] Handyside AH. 24-chromosome copy number analysis: a comparison of available technologies. Fertil Steril. 2013;100(3):595–602.

[48] Treff NR, Forman EJ, Scott RT Jr. Next-generation sequencing for preimplantation genetic diagnosis. Fertil Steril. 2013;99(6):e17–e18.

[49] Vaisvila R, Ponnaluri VKC, Sun Z, Langhorst BW, Saleh L, Guan S, et al. Enzymatic methyl sequencing detects DNA methylation at single-base resolution from picograms of DNA. Genome Res. 2021;31(7):1280–1289.

[50] Ewels PA, Peltzer A, Fillinger S, Patel H, Alneberg J, Wilm A, et al. The nf-core framework for community-curated bioinformatics pipelines. Nat Biotechnol. 2020;38(3):276–278.

[51] Saenz-de-Juano MD, Ivanova E, Romero S, Lolicato F, Sánchez F, Van Ranst H, et al. DNA methylation and mRNA expression of imprinted genes in blastocysts derived from an improved in vitro maturation method for oocytes from small antral follicles in polycystic ovary syndrome patients. Hum Reprod. 2019;34(9):1640–1649.

[52] Okae H, Chiba H, Hiura H, Hamada H, Sato A, Utsunomiya T, et al. Genome-wide analysis of DNA methylation dynamics during early human development. PLoS Genet. 2014;10(12):e1004868.

[53] De Vos M, Smitz J, Thompson JG, Gilchrist RB. The definition of IVM is clear-variations need defining. Hum Reprod. 2016;31(11):2411–2415.

[54] Lin YH, Hwang JL, Huang LW, Mu SC, Seow KM, Chung J, et al. Combination of FSH priming and hCG priming for in-vitro maturation of human oocytes. Hum Reprod. 2003;18(8):1632–1636.

[55] Pincus G, Enzmann EV. THE COMPARATIVE BEHAVIOR OF MAMMALIAN EGGS IN VIVO AND IN VITRO : I. THE ACTIVATION OF OVARIAN EGGS. J Exp Med. 1935;62(5):665–675.

